# Deciphering the comprehensive relationship between 5′ UTR and 3′ UTR sequences with deep learning

**DOI:** 10.1101/2025.05.17.654644

**Authors:** Kanta Suga, Keisuke Yamada, Michiaki Hamada

## Abstract

**Motivation:** Recent advances in mRNA therapeutics have driven further research on untranslated region (UTR) of mRNA. However, prior studies have mainly focused on either the 5^′^ or 3^′^ UTR individually. Increasing evidence suggests potential cooperative effects between these two regions, which remain largely unexplored in computational studies.

**Results:** We present a deep learning-based approach to predicting relationships between 5^′^ and 3^′^ UTRs by leveraging latent representations from a pre-trained RNA language model and contrastive learning. Our method effectively identifies highly related UTR pairs, uncovering sequence and expression characteristics that suggest functional interplay. Our analysis revealed that Highly Related UTR pairs (HRUs) are significantly enriched in genes associated with neural development, exhibit distinctive UTR length and secondary structure characteristics, and are involved in cell type-specific regulation of translation efficiency. These findings provide new insights into UTR co-optimization for mRNA therapeutics.

**Availability:** The source code is available for free at https://github.com/hmdlab/utr_pairpred.git. The data and intermediate files used in our analysis are available at https://waseda.box.com/v/utr-pairpred-data.

## 1 Introduction

Untranslated regions (UTRs) at the 5^′^ and 3^′^ ends of eukaryotic mRNAs are critical regulators of gene expression, profoundly influencing mRNA stability and translational efficiency. The 5^′^ UTR plays a crucial role in ribosome recruitment, while the 3^′^ UTR contains regulatory elements that control mRNA decay and translation. In mRNA-based therapeutics, these factors are particularly important, and a growing body of research highlights the significance of UTR sequences themselves in maximizing mRNA performance. For instance, Asrani *et al*. (2018) and Leppek et al. (2022) demonstrated that in various natural mRNAs, optimizing 5^′^-3^′^UTR combinations leads to improved translation efficiency and stability. These findings emphasize that UTR sequences are the active determinants of mRNA’s therapeutic potential.

Recent engineering efforts of UTR sequences have focused on 5^′^ or 3^′^ UTR independently (Sample *et al*., 2019; Castillo-Hair *et al*., 2024; Fu *et al*., 2024; Kirshina et al., 2023; Liu et al., 2024), while naturally occurring UTRs act in coordination to regulate translation. In a well-established closed loop model, interactions between the cap-binding complex and poly(A)-binding proteins physically bridge the 5^′^and 3^′^ ends of an mRNA, facilitating translation initiation (Tomek and Wollenhaupt, 2012; Archer *et al*., 2015; Kim *et al*., 2023). Beyond this general mechanism, certain transcripts modulate translation levels via direct base-pairing interactions between their 5^′^ and 3^′^ UTRs (Guo *et al*. (2001); Chen and Kastan (2010); Ruiz de los Mozos *et al*. (2013); Braun *et al*. (2017)). Therefore, for both basic science and engineering, it is crucial to understand the sequence features that regulate the interaction between 5^′^ and 3^′^ UTRs.

The increasing availability of large-scale genomic data has enabled us to thoroughly investigate 5^′^–3^′^UTR relationships. With the widespread adoption of high-throughput techniques, such as RNA sequencing (RNA-seq) (Wang *et al*., 2009) and ribosome profiling (Ribo-seq) (Ingolia *et al*., 2009), the expansion of large RNA sequence databases provides a valuable resource for data-driven analysis (The-RNAcentral-Consortium, 2017).

Recent advances in deep learning models for natural language processing have spurred significant research into RNA language models, representing a novel computational approach in bioinformatics (Zhang *et al*. (2024), Chu *et al*. (2024), Yu *et al*. (2024)). RNA-FM (Chen *et al*., 2022) is one such language model, trained on millions of RNA sequences (The-RNAcentral-Consortium, 2017). Embeddings generated by RNA-FM demonstrated superior predictive performance compared to conventional methods across diverse RNA-related tasks, such as mean ribosome loading prediction, protein-RNA interaction prediction and secondary structure prediction. In addition, RiNALMo (Penić *et al*., 2024), which was trained on a larger corpus of RNA sequences using a more sophisticated model architecture, is also a representative RNA language model and has demonstrated high performance across more diverse range of tasks, including splice-site prediction. This success suggests that RNA language models can capture more flexible and biologically meaningful embedding than traditional methods.

In parallel, contrastive learning (Chen *et al*., 2020) has emerged as a powerful technique for learning representations that associate information across different domains. A notable example comes from protein science, where proteins and their ligands (e.g., drugs) are embedded into a joint representation space. This protein–ligand co-embedding approach dramatically improved the prediction of molecular interactions (Singh *et al*., 2023; Wang *et al*., 2024; Zhang *et al*., 2025). Inspired by such approaches, a similar strategy can be applied to 5^′^ and 3^′^ UTRs to obtain meaningful joint representations.

Here, we propose a computational method to analyze the relationship of 5^′^ and 3^′^ UTR sequences in mRNA. By combining latent representations extracted from a pre-trained RNA language model (RNA-FM, RiNALMo) with contrastive learning (Chen *et al*., 2020), we enabled the model to distinguish UTR pairs derived from the same mRNAs from those that are not. Further analysis focusing on highly related UTR paris (HRUs) revealed that specific combinations of UTRs may play a significant role in regulating translation levels and interactions with RNA binding proteins (RBPs) involved in mRNA processing. This study presents the novel computational method for comprehensively investigating the relationship between the 5^′^ and 3^′^ UTRs, providing an avenue to elucidate the synergistic regulatory roles of UTR interactions and to guide the rational design of advanced mRNA therapeutics.

## 2 Materials and Methods

### 2.1 Data Collection

In this study, we constructed datasets of untranslated region (UTR) sequences by extracting GENCODE protein-coding transcripts (Frankish *et al*., 2022) from human (gencode.v44) and mouse (gencode.vM33) genomes. We selected transcripts that contained both 5^′^ and 3^′^ UTRs, each exceeding 10 nucleotides in length. For genes with multiple variants, only the longest transcript was retained. Redundant sequences were removed using CD-HIT (Li and Godzik, 2006) with a sequence similarity threshold of 0.9, and this filtering was applied, respectively, to 5^′^ and 3^′^ UTR sets. The final dataset comprised 16,819 transcripts for human and 18,165 for mouse.

### 2.2 Sequence embedding with RNA language model

The UTRs of endogenous mRNAs vary greatly in length, ranging from hundreds to thousands of nucleotides. Due to the length diversity, directly inputting the entire nucleotide sequence into a model is challenging. To address this challenge, we converted UTR sequences into fixed-dimensional embeddings using an RNA language model and provided these embeddings as input to the prediction model.

We employed two state-of-the-art pre-trained models designed for various RNA-related tasks: RNA-FM (hidden dimension = 640) and RiNALMo (hidden dimension = 1280). RNA-FM is a Transformer encoder-based model with 100 million parameters, trained on 23.7 million ncRNA sequences from the RNAcentral database (The-RNAcentral-Consortium, 2017). On the other hand, RiNALMo was trained on a larger dataset of 36 million RNA sequences from RNAcentral and other databases such as Rfam and Ensembl. RiNALMo has a BERT-style architecture with 650 million parameters, incorporating advanced techniques such as rotary positional embeddings (RoPE) (Su *et al*. (2024)), SwiGLU activation functions (Shazeer (2020)), and FlashAttention2 (Dao (2023)). To obtain embeddings using RNA-FM, we extracted the embedding corresponding to the [CLS] token from the model’s output for a given input sequence. For UTR sequences exceeding the model’s maximum input length (1022 nucleotides), the sequence was segmented at this length, [CLS] token embeddings were obtained for each segment, and mean pooling was applied to generate the final embedding (e.g., a sequence of 2050 nt was divided into segments of 1022, 1022, and 6 nucleotides, and embeddings from each segment were then combined through mean pooling). In contrast to RNA-FM, RiNALMo employs relative positional embeddings, and it does not have a maximum input length constraint. Therefore, we obtained a *D*-dimensional embedding from RiNALMo by mean pooling the *L × D* output embedding matrix along the *L* dimension, where *L* represents sequence length.

### 2.3 Data Pair Construction for Prediction Tasks

For our prediction task, we constructed positive and negative UTR pairs as illustrated in Fig. 1a. Positive pairs consisted of 5^′^ UTR and 3^′^ UTR sequences derived from the same mRNA transcript, representing naturally co-occurring UTR combinations. Negative pairs were created by pairing 5^′^ UTR sequences with 3^′^ UTR sequences from different mRNA transcripts, representing non-native combinations. This paired dataset construction allowed us to train models that could distinguish between biologically relevant and random UTR associations.

**Fig. 1.**
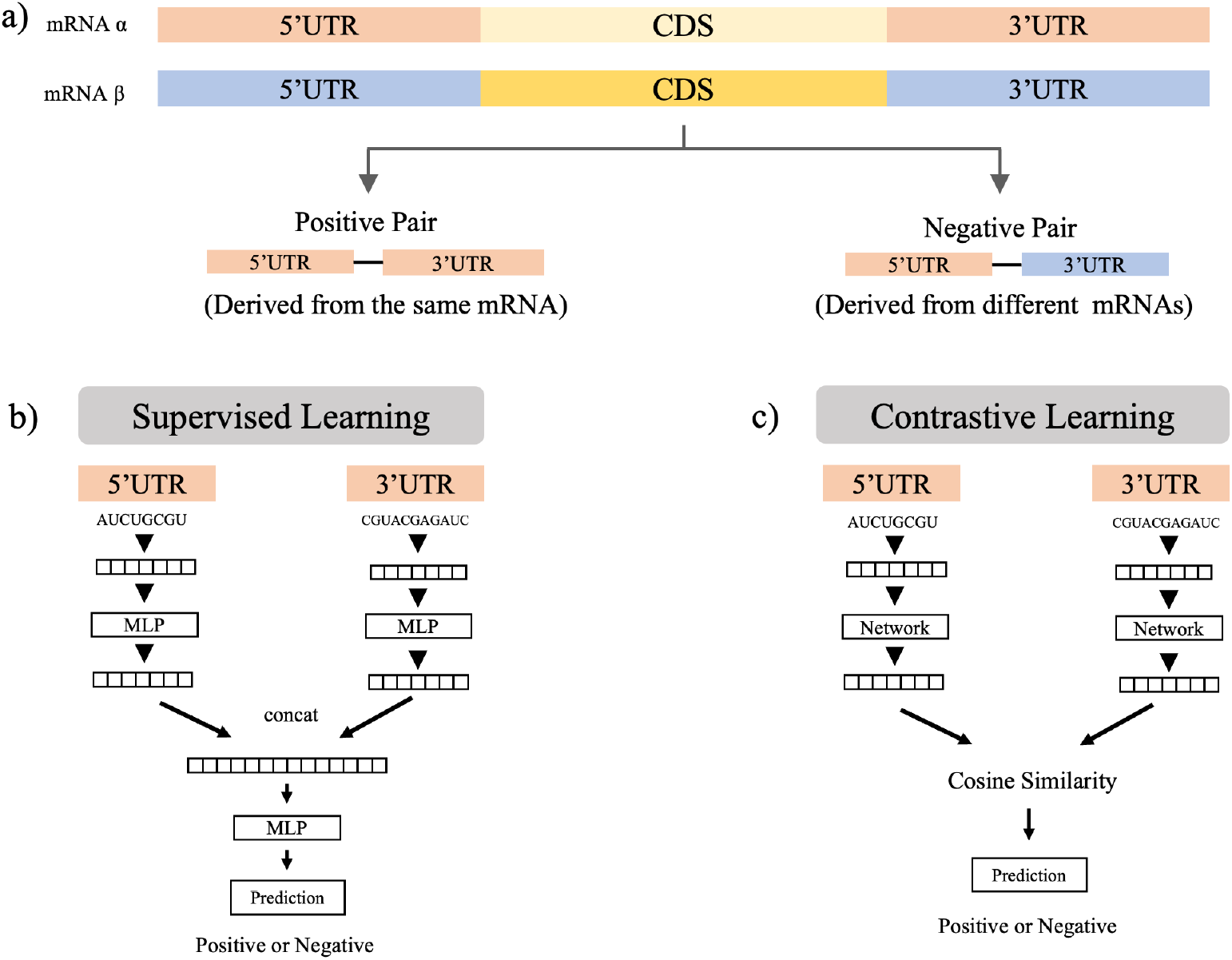
**a)** Data construction for the UTR pair prediction task. Pairs of 5^*′*^UTR and 3^*′*^UTR sequences derived from the same mRNA were designated as positive pairs, while those from different mRNAs were defined as negative pairs. The prediction task aims to classify a given UTR pair as positive or negative with either **b)** Supervised learning or **c)** Contrastive learning method. In both methods, sequence embeddings extracted from a pre-trained RNA language model were used as the inputs for the prediction task.

### 2.4 Training Methods

We employed two independent approaches–supervised learning and contrastive learning–to predict the relationship between 5^′^ and 3^′^ UTRs.

#### 2.4.1 Supervised learning

During supervised learning, the model was trained using explicitly labeled data (Fig. 1b). The 5^′^ and 3^′^ UTR sequences were passed through separate multilayer perceptron (MLP) layers, and their outputs were concatenated into a single vector. This concatenated vector was then reduced in dimensionality through three additional MLP layers, producing logits for prediction.

A target label of 1 was assigned to the pairs of 5^′^ and 3^′^ UTRs originating from the same mRNA, and 0 otherwise. The model was trained using binary cross-entropy loss to learn the correspondence between UTR pairs and their labels. After hyperparameter tuning, we set the batch size to 32, the learning rate to 1e-4, and used the Adam optimizer. To mitigate overfitting, we applied a dropout rate of 0.5.

#### 2.4.2 Contrastive learning

We also adapted a contrastive learning approach (Chen *et al*., 2020) to predict UTR pair relationships. In this approach, multiple positive pairs of 5^′^ UTR and 3^′^ UTR sequences from the same mRNA were collected into a batch. Within a batch, the model was trained to bring the representations of positive 5^′^–3^′^ UTR pairs closer together while pushing apart the representations of non-matching UTR pairs.

In contrastive learning, the similarity between UTR pairs was measured using the cosine similarity between their vector representations. During testing, we calculated the cosine similarity for a given 5^′^–3^′^ UTR pair and applied a similarity threshold (e.g., 0.0) to determine whether the pair originated from the same mRNA.

We treated the cosine similarity between the representations of 5^′^ and 3^′^ UTRs from contrastive learning as a quantitative measure of their relation score. By performing 10-fold cross-validation, we computed a pseudo-relatedness score for all mRNAs in the dataset. Hereafter, we refer to this cosine similarity as the relation score of the UTR pair (Figure 1(c)).

#### 2.4.3 Random forest

As a baseline for performance comparison with the proposed method, we employed a random forest classifier (Breiman, 2001). Similarly, the random forest model predicted whether a given pair of 5^′^ and 3^′^ UTR sequences originated from the same mRNA. Following the approach of Cao et al. (2021), we extracted various features from the sequences, such as length, GC content, energy, and k-mer frequency, and provided a total of 5,450 features as input to the model.

### 2.5 Analysis for highly related UTRs

To elucidate biological factors contributing to the highly related UTRs, we conducted analysis from multiple perspectives, involving with the sequence features of UTRs (2.5.1), mRNA-RBP interactions (2.5.2) and translation regulation (2.5.3).

#### 2.5.1 Length & MFE comparison

We performed a comparative analysis of the minimum free energy (MFE). MFE is an important indicator for the structural functionality of RNA sequence. As illustrated in Fig. 3a, the UTR regions were divided into subclasses for detailed comparison. In the case of the 5^′^ UTR, we compared the first 100 nucleotides (designated as 5^′^ UTR head) and the last 100 nucleotides (5^′^ UTR tail), as well as a 200-nucleotide region comprising the last 100 nucleotides of the 5^′^ UTR and the first 100 nucleotides of the CDS (designated as 5^′^ UTR_tail_cds). Because the 3^′^ UTR sequences are generally longer, we calculated the MFE for the first and last 500 nucleotides (designated as 3^′^ UTR head and 3^′^ UTR tail, respectively) and for a 600-nucleotide region formed by the junction of the last 100 nucleotides of the CDS with the beginning of the 3^′^ UTR (designated as 3^′^ UTR_head_cds). MFE calculations were performed using RNAfold Lorenz *et al*. (2011).

#### 2.5.2 Features of RBP interactions

To investigate the association between HRUs and mRNA-RBP interactions, we conducted comprehensive analysis using eCLIP (Van Nostrand *et al*., 2016) data. eCLIP is a transcriptome-wide sequencing method to map RNA regions that interact with RBPs. On the mRNA side, we categorized transcripts into groups based on relation score thresholds, following the approach shown in Figure 2(a). On the RBP side, we collected all available human eCLIP BED files from the ENCODE database (Moore *et al*., 2020). We calculated the number of eCLIP peak intersecting with the UTRs using bedtool (Quinlan and Hall, 2010). Subsequently, the mean number of peaks was computed within each UTR relation score group. These scores were individually caluculated for 5^′^ and 3^′^ UTRs. To identify RBPs that preferentially bind to HRUs, we computed the ratio of the average intersection values between the HRUs (0.7-1.0) and the lowest relation score group (0.0–0.1). We then focused on the top 30 RBPs with the highest ratios for the downstream gene ontology analysis.

**Fig. 2.**
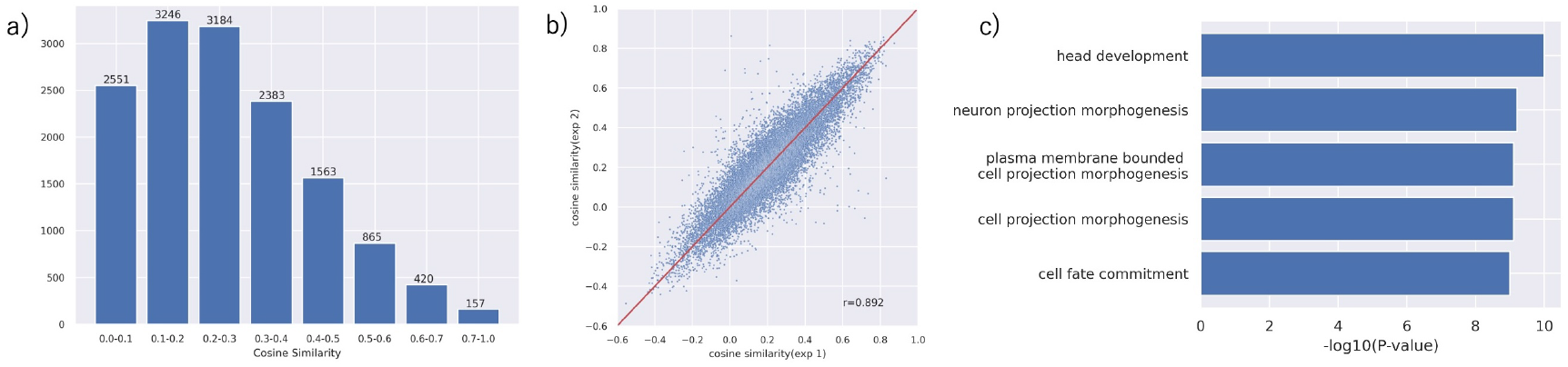
(a) Distribution of cosine similarity values for pairs of 5^*′*^ and 3^*′*^ UTRs of human mRNAs obtained using contrastive learning combined with RiNALMo embedding. The subset with the highest similarity range (0.7∼1.0; n = 157) was defined as “Highly Related UTRs (HRUs)” for further analysis. (b) Cosine similarity results obtained by repeating the contrastive learning + RiNALMo approach with different random seeds. The strong correlation observed across experiments indicates that the results used for the analysis are not coincidental. (c) The top five Gene Ontology terms with the highest −log10(p value) from the gene enrichment analysis of HRUs.

**Fig. 3.**
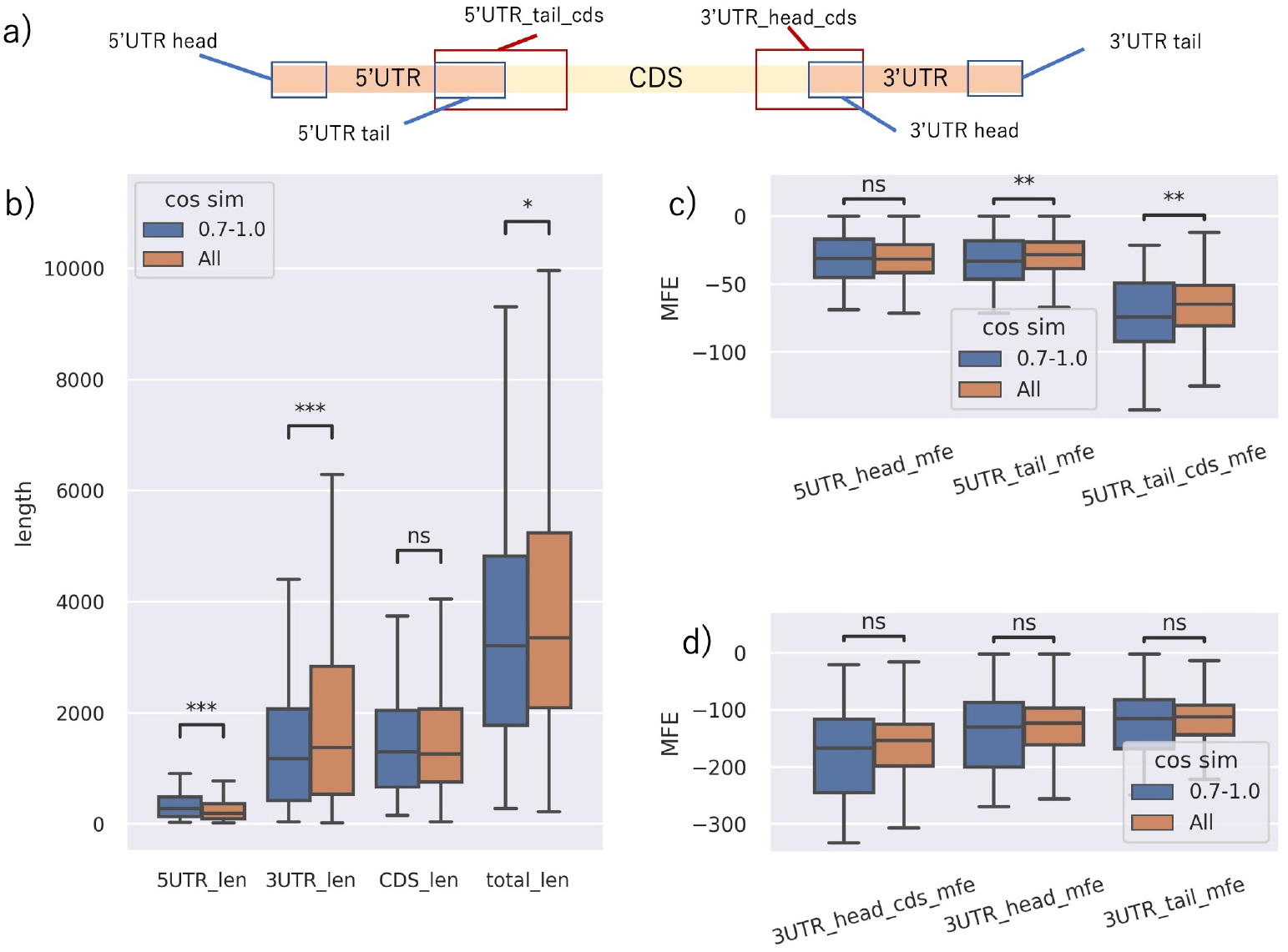
Sequence feature analysis. a) The names of the subdivisions for each analyzed mRNA region. Comparison of each mRNA region between HRUs and all other transcripts about b) sequence length, c) MFE (kcal/mol) in each subdivision of the 5^′^UTR, d) MFE in each subdivision of the 3^′^UTR.

#### 2.5.3 Translation efficiency

To assess differences in translation efficiency between HRUs and the all other mRNA, we calculated translation efficiency (TE) across multiple human cell types and tissues. TE was calculated as

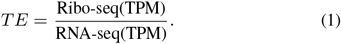

following the approach by Chothani *et al*. (2019). Raw RNA-seq and Ribo-seq reads were obtained from The Sequence Read Archive (Leinonen *et al*., 2010) and mapped using bowtie2 (Langmead and Salzberg, 2012). The detailed data information is provided in Supplementary Table S1. For statistical analysis, the Mann-Whitney U test was used to assess differences in TE distributions between HRUs and all other transcripts.

## 3 Results

### 3.1 Performance comparison

For the task of predicting the correspondence between the 5^′^ and 3^′^ UTR on the same mRNA, we evaluated four methods, each combining an RNA language model (RNA-FM or RiNALMo) with a different learning strategy (supervised learning or contrastive learning). Additionally, as a baseline, we implemented a random forest classifier with extracted sequence features and compared its performance. The results are presented in Table 1. RNA language model-based methods (RNA-FM and RiNALMo) significantly outperformed the random forest baseline. Both models showed enhanced performance with contrastive learning compared to supervised learning. Notably, RiNALMo, likely benefiting from its deeper architecture, higher-dimensional embeddings, consistently achieved the highest scores. These results suggest that contrastive learning with a large-scale RNA model is a promising approach for estimating relationships between 5^′^ and 3^′^ UTRs.

**Table 1.**
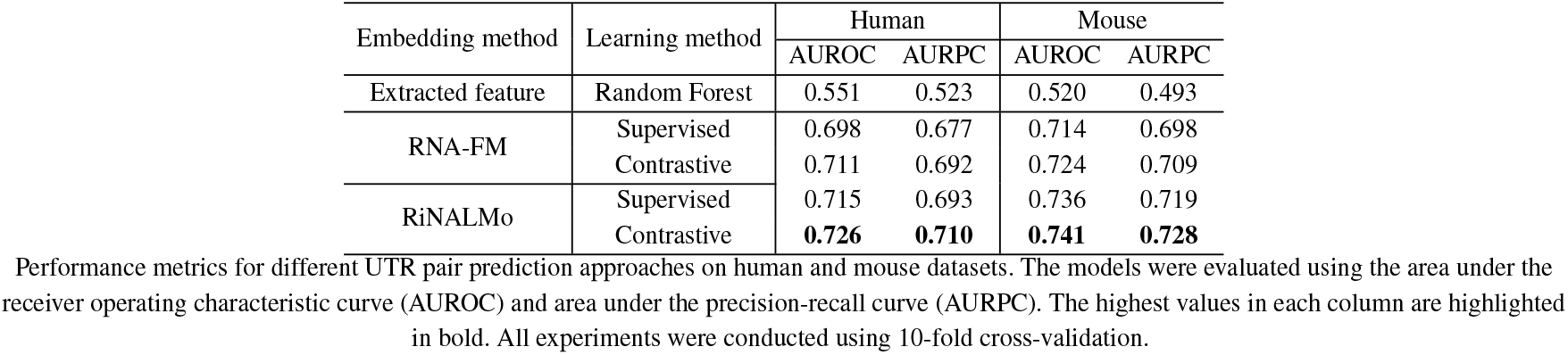
Performance comparison of UTR pair prediction methods across different embedding and learning approaches.

### 3.2 Analysis for highly related UTRs

We investigated the factors contributing to the relationship between untranslated regions (UTRs) on the same mRNA from multiple perspectives. For our analysis, we utilized the results obtained from the combination of contrastive learning and RiNALMo, which achieved the highest performance in the UTR pair prediction task. In the model trained with contrastive learning, cosine similarity was computed for each input pair of 5^′^ and 3^′^ UTRs. We defined this cosine similarity as a relation score between the two regions. A higher relation score may suggests a stronger functional association between the 5^′^ and 3^′^ UTR, indicating that their co-occurrence on the same mRNA is biologically significant. Using 10-fold cross-validation, we obtained relation scores for all mRNAs in the human dataset; the distribution of these scores is shown in Figure 2(a). Subsequent analyses focused on samples with the highest relation scores (0.7–1.0, n = 157), which we designated as **H**ighly **R**elated **U**TR**s** (HRUs). When we calculated the relation scores for all mRNAs in two independent experiments, a strong correlation was observed between experiments, confirming the reproducibility of these scores (Fig. 2b).

To investigate whether the genes associated with HRUs exhibit any distinctive characteristics, we performed a gene enrichment analysis using Metascape (Zhou *et al*. (2019)). The results revealed a significant enrichment of genes involved in developmental processes, particularly “head development” and “neuron projection morphogenesis”, suggesting that interactions between UTRs are involved in the translation regulation of those genes (Fig. 2c).

#### 3.2.1 Analysis for sequence features

We analyzed the sequence characteristics of HRU-containig mRNAs by dividing each UTR region into detailed subregions and analyzing their features (Fig. 3a). First, we compared the lengths of each region between HRU-containig mRNAs and all other mRNAs. This analysis revealed that, relative to the overall mRNA population, HRU-containig mRNAs exhibit a significantly longer 5^′^ UTR, a significantly shorter 3^′^ UTR, no significant difference in CDS length, and an overall significantly shorter transcript length (Fig. 3b).

To investigate whether HRUs exhibit distinctive structural characteristics, we compared the MFE values between mRNAs with HRUs and all other mRNAs in each subregions. The mRNAs with HRUs exhibited significantly lower MFE value at the end of 5^′^ UTR and at the intersection between 5^′^ UTR and CDS, suggesting that secondary structure formation in 5^′^UTR is more pronounced. Collectively, mRNAs with HRUs tend to have shorter untranslated regions and more stable secondary structures. These sequence and structural features are likely to influence various aspects of gene expression regulation, including mRNA stability and translation efficiency.

#### 3.2.2 Analysis for interactions with RBPs

RNA binding proteins (RBPs) are known to significantly influence RNA function through interactions with UTRs (Liu and Cao, 2023). To investigate RBP-binding profiles in HRUs and other UTR pairs, we performed a cross-analysis between the UTRs and eCLIP data. Clustering analysis revealed a distinct group of RBPs that preferentially bind to HRUs in both the 5^′^ and 3^′^ UTRs (Fig. 4ab). Subsequent gene enrichment analysis of these RBPs indicated significant enrichment in genes associated with mRNA metabolic processes and mRNA processing, suggesting a strong functional relevance of these interactions (Fig. 4cd).

**Fig. 4.**
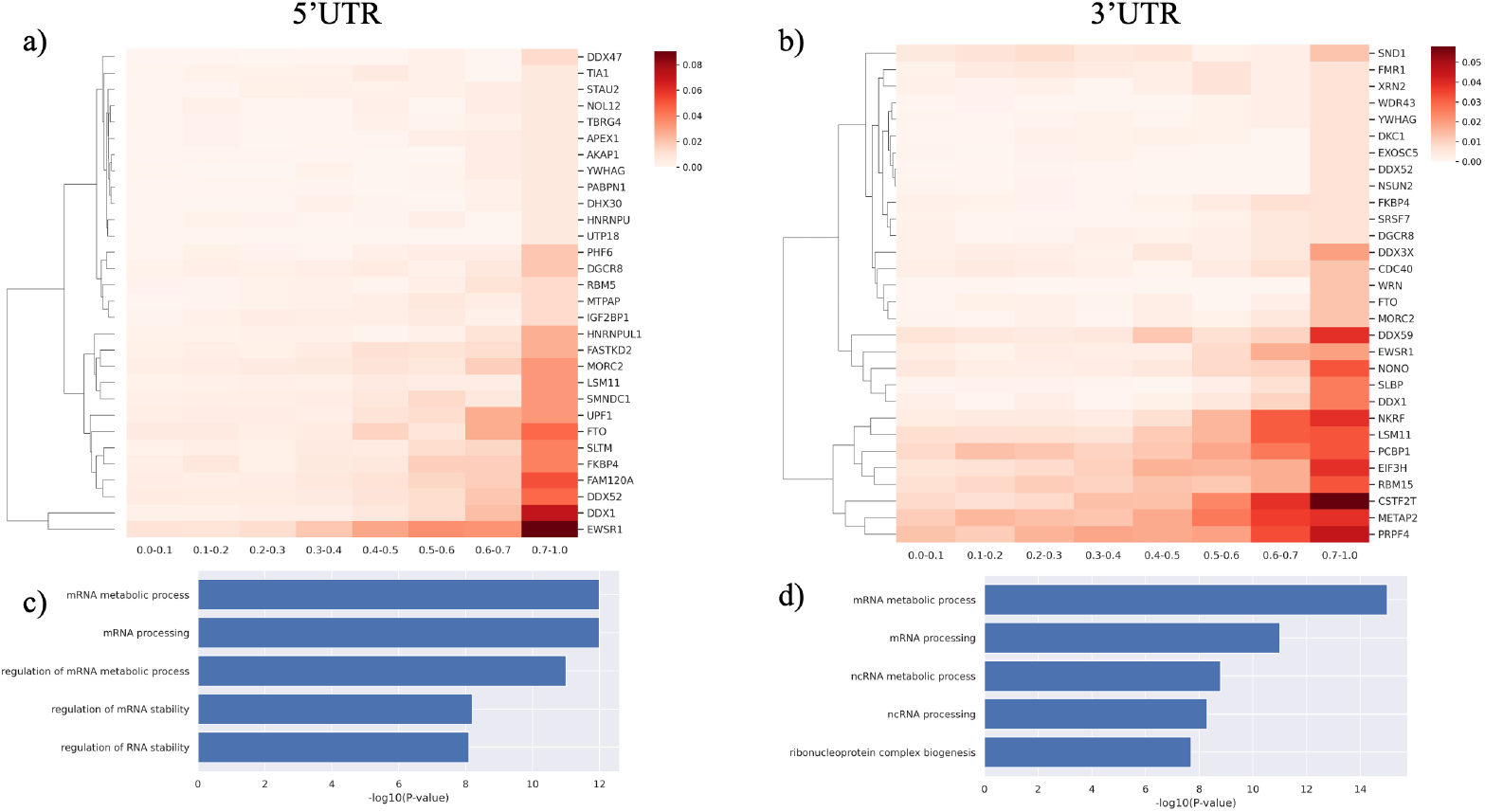
Analysis of interactions between 5^*′*^/3^*′*^ UTRs and RNA-binding proteins (RBPs). The intersection between all eCLIP datasets in ENCODE and the UTR regions was first calculated and averaged for each threshold of the relation score. By comparing the mean values of HRUs with those of the 0.0–0.1 relation score group, the top 30 RBPs were identified. Panels (a) and (b) show cluster heatmaps for these 30 RBPs in the 5^*′*^ and 3^*′*^ UTR, respectively, while panels (c) and (d) present the results of functional enrichment analysis for the 5^*′*^ and 3^*′*^ UTR, respectively.

#### 3.2.3 Analysis for translation efficiency

The UTR sequence significantly regulates the mRNA translation levels. To investigate whether substantial differences in translation efficiency (TE) exist between HRUs and other mRNAs, we calculated and compared TEs across multiple cell types and tissues. Interestingly, as shown in Fig. 5, mRNAs containing HRUs exhibited significantly higher TE relative to the overall average in certain cell types (HepG2: *P* = 3.57 × 10^−6^, HEK293: *P* = 4.98 × 10^−3^), whereas they displayed significantly lower TE in others (Breast:*P* = 2.09 × 10^−7^, PC3:*P* = 3.21 × 10^−4^, HeLa:*P* = 3.39 × 10^−4^). In some other cell types, there was no significant difference in TE (Supplementary Figure S1). These findings suggest that HRUs may contribute to the regulation of translation efficiency in a cell-line or tissue specific manner.

**Fig. 5.**
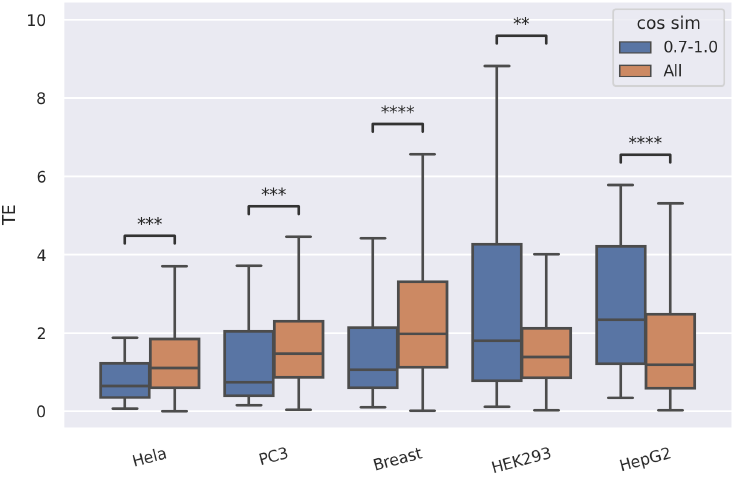
Comparison of translation efficiency (TE) between HRUs and other mRNAs across different cell lines by the Mann-Whitney U test. TE was analyzed separately for each cell line. In HepG2 and HEK293 cells, HRUs exhibited significantly higher TE, whereas in HeLa, PC3, and Breast cell lines, HRUs showed significantly lower TE.

## 4 Discussion

In this study, we investigated predictable relationships between the 5^′^ and 3^′^ UTRs within the same mRNA molecule. To explore this, we employed a deep learning-based approach that integrated sequence prediction with multi-faceted downstream analyses. In our paired prediction task, incorporating sequence embeddings from the RNA pre-training model RiNALMo with contrastive learning achieved the highest prediction accuracy. The strong performance of our predictive model suggests a non-negligible association between the 5^′^ UTR and 3^′^ UTR sequences.

Leveraging association scores from our prediction task, subsequent downstream analyses further revealed that mRNAs with high UTR relational scores (HRUs) possess distinct sequence characteristics in terms of both length and MFE, along with different expression profiles compared to other mRNA populations. Notably, HRUs were enriched for transcripts encoding genes involved in brain development and neural function. HRU-containing mRNAs also exhibited a pronounced tendency to interact with RNA-binding proteins (RBPs) implicated in mRNA processing and showed significant variability in translation efficiency across cell types.

Although our findings indicate a clear association between 5^′^ and 3^′^ UTRs, the underlying molecular mechanisms remain elusive. We found that certain RBPs, which preferentially bind to HRUs, have gene ontologies commonly associated with mRNA processing in both 5^′^ and 3^′^ UTRs. This finding suggests that the interplay between UTRs may be involved in recruiting RBPs that mediate mRNA processing. For instance, in human p53 mRNA, RNPC1 is known to bind to both 5^′^ and 3^′^ UTRs, thereby inhibiting the association of the translation initiation factor eIF4E (Zhang *et al*., 2011). A similar mechanism may also occur in the HRUs we have found in this study.

Future investigations of the mechanistic basis of 5^′^–3^′^ UTR interactions may uncover novel regulatory pathways in mRNA translation. A deeper understanding of these processes could not only enhance our knowledge of post-transcriptional regulation but also contribute to the rational design of next-generation mRNA therapeutics.

## Author Contributions

K.S. and M.H. designed and conceived this study. K.S. performed computational experiments and analyses. K.Y. and M.H. supervised the project. All authors contributed to the writing and review of the manuscript.

## Acknowledgements

We would like to express our sincere appreciation to the members of the Hamada Laboratory, especially Dr. Masato Kosuge. We are also grateful for the insightful discussions with members of the AMED/LEAP project, particularly Dr. Hiroshi Abe, Dr. Satoshi Uchida and Dr. Yoshihiro Shimizu.

## Funding

This work was supported by AMED under Grant Number 24gm0010008 to M.H.

## Notes

### Competing Interest Statement

The authors have declared no competing interest.

