## Supplemental data for "Deciphering the comprehensive relationship between 5′ UTR and 3′ UTR sequences with deep learning"

### 1 Supplementary Figures and Tables

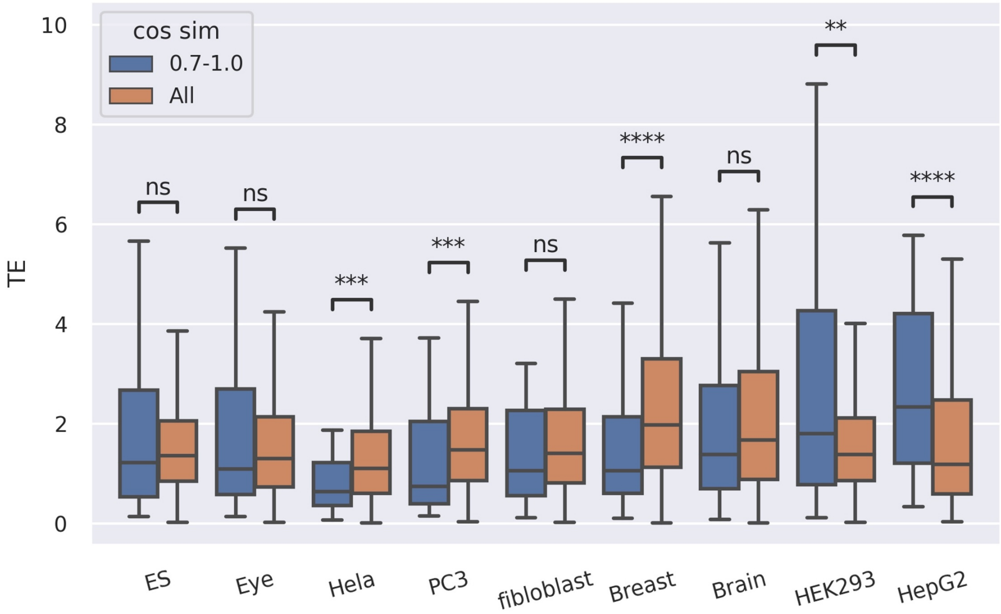


Fig. S1: Comparison of translation efficiency (TE) between HRUs and other mRNAs across 9 kinds of cell lines/tissue by the Mann-Whitney U test. TE was analyzed separately for each cell line/tissue

Table S1: A list of SRR-ID used for TE comparison analysis

| SRR ID | Cell line/Tissue | Sequence type | Reference |
|---|---|---|---|
| SRR1562539 | Brain | ribo-seq | Gonzalez *et al.* (2014) |
| SRR1562544 | Brain | rna-seq | |
| SRR2075925 | HEK293 | ribo-seq | Iwasaki *et al.* (2016) |
| SRR2075930 | HEK293 | rna-seq | |
| SRR057526 | Hela | ribo-seq | Guo *et al.* (2010) |
| SRR057527 | Hela | rna-seq | |
| SRR403883 | PC3 | ribo-seq | Hsieh *et al.* (2012) |
| SRR403882 | PC3 | rna-seq | |
| SRR2064024 | fibloblast | rna-seq | Tirosh *et al.* (2015) |
| SRR2064017 | fibloblast | ribo-seq | |
| SRR1204658 | Muscle | rna-seq | Wein *et al.* (2014) |
| SRR1204656 | Muscle | ribo-seq | |
| SRR1610260 | ES | rna-seq | Werner *et al.* (2015) |
| SRR1610244 | ES | ribo-seq | |
| SRR1573941 | Breast | rna-seq | Rubio *et al.* (2014) |
| SRR1573939 | Breast | ribo-seq | |
| SRR1976453 | Eye | rna-seq | Tanenbaum *et al.* (2015) |
| SRR1976447 | Eye | ribo-seq | |
| SRR8135336 | HepG2 | ribo-seq | Huang *et al.* (2019) |
| SRR8131644 | HepG2 | rna-seq | |
